# Eyes wide open: regulation of arousal by temporal expectations

**DOI:** 10.1101/2020.07.24.219428

**Authors:** Nir Shalev, Anna C. Nobre

## Abstract

To perform a task continuously over an extended period of time, it is necessary to maintain an adequate level of arousal. In cognitive research, traditional studies have used repetitive, monotonous tasks to learn about the dynamics of arousal in tasks that require sustained vigilance, such as driving or monitoring a factory line. To date, studies have rarely considered whether observers use task-embedded regularities in such continuous contexts to anticipate events and regulate arousal accordingly. In the current study, we explored whether arousal is an adaptive process that reacts to temporal stimulus predictability. Across two experiments, we used pupillometry as a proxy measure of arousal in human observers performing continuous tasks. Within the tasks, we manipulated the temporal predictability of relevant events by presenting stimuli either in a fixed rhythm or at varied intervals. Temporal predictability led to the lowering of tonic levels of arousal. Trial-wise temporal structures led to short-scale changes in pupil size related to temporal anticipation. Accordingly, we suggest that arousal is sensitive to varying levels of temporal predictability and dynamically adapts to task demands to bring about performance benefits as well as energetic efficiencies overall.

**Statement of Relevance:** People often have to sustain focus and high levels of performance on extended and non-stimulating tasks (e.g., driving, sowing, monitoring data acquisition). A critical factor to the success (or failure) in sustained performance is arousal – the ‘energetic state’ of the cognitive system. Here we used pupil dilation as a proxy to study levels of arousal during sustained performance on monotonous tasks. We reveal that arousal is dynamically regulated to support performance according to an important fundamental property of any task: its temporal structure. When the timing of task-relevant events is predictable, arousal levels fluctuate accordingly, saving energy overall while also optimally guiding performance. Our study and findings add ecological validity to the study of temporal expectations, by moving investigations beyond typical trial-by-trial designs. They also carry significant implications for clinical studies relying on sustained performance tasks.

## Introduction

The ability to anticipate the timing of an upcoming event allows us to prepare and perform more efficiently (Nobre & Van Ede, 2018). In experimental settings, temporal expectations are typically manipulated on a trial-by-trial basis, to estimate the behavioural benefits conferred by temporally informative cues (J T Coull & Nobre, 1998), short rhythmic stimulations (Jones, 2010), learned temporal probabilities (Ghose & Maunsell, 2002), or sequential effects (Capizzi, Correa, Wojtowicz, & Rafal, 2015; Los, 2010). These experimental approaches are highly relevant when trying to map the mechanisms that support temporal anticipation of discrete cognitive events. Much less is known about how anticipation of events unfolds when we engage continuously with extended cognitive tasks. Arguably, the transition from discrete task events towards continuous task performance is an important step toward understanding human behaviour in natural settings. Many forms of human behaviour require sustained focus and repetitive patterns, such as driving home from work or sowing a quilt blanket or monitoring a data stream during an experimental session for occasional artifacts. In such contexts, behaviour relies heavily on the capacity to maintain an adequate state of arousal over an extended period of time, and is not fully captured by mechanisms restricted to the momentary engagement with perceptual events (Shalev, Bauer, & Nobre, 2019).

Experimentally, sustained performance is typically studied using Continuous-Performance Tasks (CPT). Most CPT studies have emphasised the role of slow modulations in sustained attention and arousal functions as the key factor in performance (Davies & Parasuraman, 1982; Robertson & O’Connell, 2010; Sarter, Givens, & Bruno, 2001). The modulation of arousal is typically attributed to the passive, gradual habituation related to repetitive stimulation (Fortenbaugh, DeGutis, & Esterman, 2017; Mackworth, 1968; Parasuraman, Warm, & Dember, 1987). However, it is unknown whether briefer, embedded task-dependent regularities affording selective temporal expectations can also affect arousal.

In CPT, the temporal structure is typically defined by the intervals between successive perceptual events, irrespective of motor responses. For example, imagine yourself practising your tennis skills using a tennis-ball machine. The machine continuously shoots tennis balls at a certain pace, regardless of whether you managed to hit a ball. The machine may shoot balls at regular or irregular intervals. In principle, if the observer is sensitive to the temporal structure of events, then two factors likely influence performance: sustaining an appropriate state of arousal and selectively deploying temporal attention to an imperative stimulus (i.e., the tennis ball). In practice, little is known about the potential of such rhythm-induced selective temporal anticipation to punctuate the overall background state of arousal.

Hebb (1955) discussed arousal as a non-specific cognitive resource that “in effect makes organised cortical activity possible”. The effect of arousal on performance is characterised by an inverted U-shaped curve: with performance best at intermediate arousal levels and falling off progressively toward the two extremes of sleep (low arousal) and anxiety (high arousal) (Aston-Jones & Cohen, 2005; Hebb, 1955). Neural control of arousal occurs through noradrenergic modulation arising in locus coeruleus of the brainstem reticular activating system (Steriade, 1996), which in turn is regulated by cortical feedback (Aston-Jones & Cohen, 2005). In humans and in non-human primates, the firing rate at the locus coeruleus-noradrenergic system and associated arousal levels are marked by changes to pupil size (Joshi & Gold, 2020; Joshi, Li, Kalwani, & Gold, 2016; Rajkowski, 1993). Thus, under the right control conditions, pupil size provides a convenient indirect proxy measure for the arousal-related system.

The ability of predictable temporal structures in extended tasks to drive transient fluctuations in arousal is particularly important given wide utilisation of CPT tasks to characterise sustained attention in clinical populations (Lee & Park, 2006; Richards, Samuels, Turnure, & Ysseldyke, 1990; Shalev, Humphreys, & Demeyere, 2016; Shalev, Steele, et al., 2019; Sims & Lonigan, 2012; Tsal, Shalev, & Mevorach, 2005). The possible contribution of selective temporal expectations to performance during CPT was suggested by a recent study in which temporal predictability was manipulated during a CPT (Dankner, Shalev, Carrasco, & Yuval-Greenberg, 2017). Individuals monitored a stimulus stream in a continuous task and responded to occasional pre-defined targets. The results showed that when targets appeared in a fixed rhythm, participants were faster and were more likely to inhibit their eye movements before stimulus onset. To our knowledge, this study provided the first evidence for the formation and benefits of temporal expectations while performing a CPT. In our current study, we take the next important step and investigate whether performance benefits of temporal expectations within a CPT are associated with modulation of arousal levels.

Valid temporal expectations about the onsets of events in CPT can be a strong source of uncertainty reduction. Studies investigating the effects of uncertainty about stimulus identity or reward associations in the context or reinforcement learning have shown strong modulation of pupil size attributed to changes in arousal (e.g., (Friedman, Hakerem, Sutton, & Fleiss, 1973; Lavín, San Martín, Rosales Jubal, Martín, & Jubal, 2014; Preuschoff, ‘t Hart, Einhäuser, & Einhauser, 2011; Urai, Braun, & Donner, 2017; Vincent, Parr, Benrimoh, & Friston, 2019). Accordingly, we hypothesised that reduction of *temporal* uncertainty, arising from predictable rhythmic stimulus presentation within the context of CPT, would also lead to changes in pupil size. We hypothesised that increased temporal uncertainty would be associated with higher levels of tonic arousal, and that the availability of temporal regularities would enable a more efficient management of arousal.

Any CPT design inherently contains task-relevant, predictive temporal structures that can, in principle, aid performance. Even when the intervals between task-relevant stimuli vary, informative temporal structures come about through learned associations based on the trial history of temporal intervals or the cumulative conditional probability of events occurring given the span of intervals (Los, Kruijne, & Meeter, 2017; Nobre, Correa, & Coull, 2007). In addition, many CPT tasks use constant intervals and thereby provide a deterministic rhythmic structure that could support significant reduction of tonic arousal by strong temporal expectation.

Across two experiments, we measured perceptual discrimination and pupil size during continuous-performance tasks to test whether temporal expectations improve behaviour and modulate arousal. The two experiments used slightly different designs but asked the same questions, thereby ascertaining whether the findings were reproducible and generalisable. In both experiments, we manipulated whether target events occurred in a temporally predictable or unpredictable fashion. Behavioural measures of perceptual discrimination tested whether performance improved when target onsets were fully predictable. Pupil traces probed whether the overall level of temporal expectations modulated the tonic arousal tone, and tested whether local changes in temporal expectation within the task were linked to short bursts of arousal related to stimulus anticipation.

## Results

In the first experiment, individuals monitored a continuous stream of task-relevant stimuli presented either at fixed (every 3500 ms) or variable intervals (in a range between 2500 and 4500 ms). The rhythmic and variable interval conditions alternated every three minutes while participants completed a single run of the task lasting approximately 12 minutes with no breaks. Target and distractor stimuli briefly replaced a visual mask otherwise continuously presented during the inter-stimulus interval. Participants were instructed to respond when a pre-defined target appeared within the stimulus stream (66% of the stimuli were targets), while ignoring nontargets (see figure 1 and Methods section). In our second experiment, we aimed to replicate our findings. This time, participants made fine perceptual discriminations in a continuous task. They indicated the direction of an arrow that briefly replaced a mask for varying durations, in a way that allowed us to model the data using a computational model based on the theory of visual attention (TVA; (Bundesen, 1990)). Fixed and Variable intervals were presented in alternating blocks (see figure 3 and Methods section).

**Figure 1:**
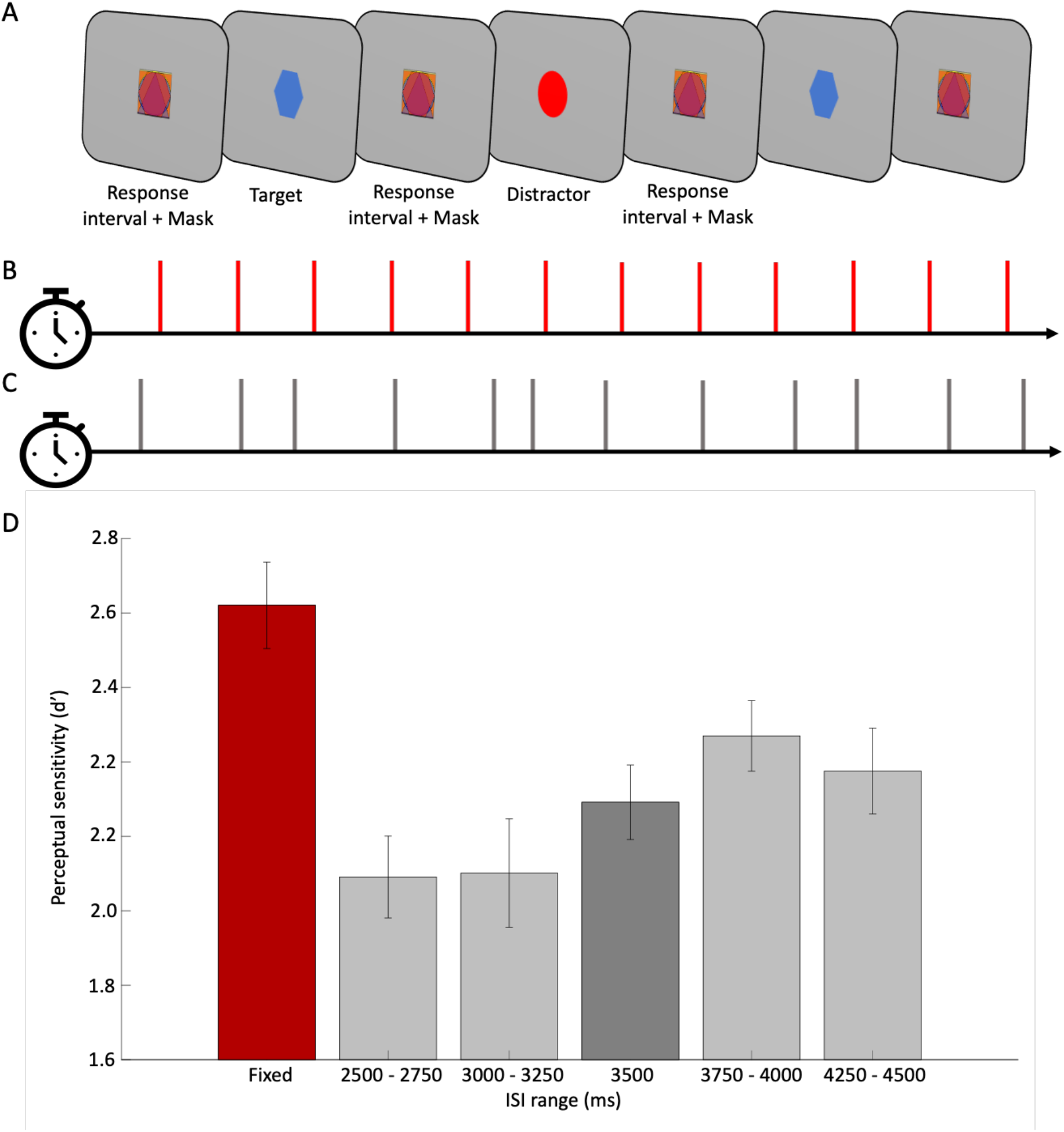
(A) a schematic illustration of the different conditions in the CPT. Participants had to respond when identifying a target (blue hexagon, designated as a ‘target’) and ignore all other shapes (for example a red circle, designated as a ‘distractor’). The stimuli – targets and distractors – were masked during the interstimulus intervals (‘mask’). Targets and distractors were presented in two conditions (blocked), which repeated two times within each experimental run. They were either presented at (B) fixed, 3500-ms intervals; or (C) at varied intervals between 2500 and 4500 ms in steps of 250 ms. In (D) we compare the perceptual sensitivity to targets appearing after 3500 ms intervals in the two experimental conditions. The red bar represents the mean perceptual sensitivity in Fixed blocks, during which all intervals were equal to 3500 ms. The grey bars depict the mean d’ on different subsets trials in variable blocks, based on the inter-stimulus intervals. Error bars represent the standard error of the mean

### Experiment 1

#### Behavioural data

Our results indicated that participants were better at detecting a target after 3500 ms when it appeared within a Fixed block, compared to when it appeared after the same interval in a Variable block (t(24)=4.05; *p*<.001 95%CI=[.259;. 798]). There were no differences in the response bias (β) (*p*=.455). Descriptive data across intervals and conditions appear in Table #1. An illustration of the task, along with the perceptual sensitivity for targets in the two conditions, appears in Figure 1.

**Table 1:** descriptive statistics. Mean and standard errors (in parenthesis) of perceptual sensitivity (d’) and response bias (β) at different intervals and conditions. The left column represents the means during Fixed blocks, when all intervals were 3500 ms. The other five columns show the average perceptual parameters in Variable blocks split by interval range: when intervals were 1) 2500 and 2750 ms; 2) 3000 and 3250 ms; 3) 3500 ms; 4) 4000 and 4250 ms; 7) 4250 and 4500 ms

|  | Fixed | Variable |  |  |  |  |
| --- | --- | --- | --- | --- | --- | --- |
|  | 3500 ms | 2500 –<br>2750 ms | 3000 –<br>3250 ms | 3500 ms | 3750 –<br>4000 ms | 4250 –<br>4500 ms |
| $d'$ | 2.82 (.11) | 2.09 (.11) | 2.10 (.14) | 2.29 (.10) | 2.46 (.09) | 2.37 (.11) |
| $\beta$ | -.16 (.08) | .03 (.06) | -.08 (.07) | -.11 (.06) | -.07 (.06) | -.09 (.07) |

In addition to the performance benefit in the fixed-interval condition, the data in Figure 1 revealed a pattern of increase in perceptual sensitivity as a function of the inter-stimulus interval in the variable-interval condition. To evaluate this pattern statistically, we used an ANOVA to test for a linear contrast. The dependant variable was the mean perceptual sensitivity, and the interval range was the independent factor. We included only trials in Variable blocks. The result indicated a significant linear trend (F(1,24)=9.668; p=.005; partial η^2^=.287). Such a pattern is in line with previous reports of the increase in the conditional probability for targets to appear as a source that drives temporal anticipation (‘hazard rates’) (Correa & Nobre, 2008; Ghose & Maunsell, 2002; Janssen & Shadlen, 2005), however in our task this could not strictly be separated from putative effects of general mounting preparation over time (Coull, 2014; Nobre, 2010).

#### Pupillometry data

Our first analysis of the pupillometry data compared tonic differences in arousal level across all trials in a block, as a marker of adapting arousal to the temporal properties of the task. We contrasted the mean pupil size over the full duration of Fixed vs. Variable blocks. Figure 2a shows the average standardised pupil size separated by block type. The two standardised signals were compared using a paired-sample t-test with the mean standardised pupil size in each condition as the dependent variable (see Figure 2b). There was a significant difference (t(24)=-2.88; p=.008; 95%CI[-1.00; −.16]), with pupil size significantly larger over the Variable block compared to the Fixed block.

**Figure 2:**
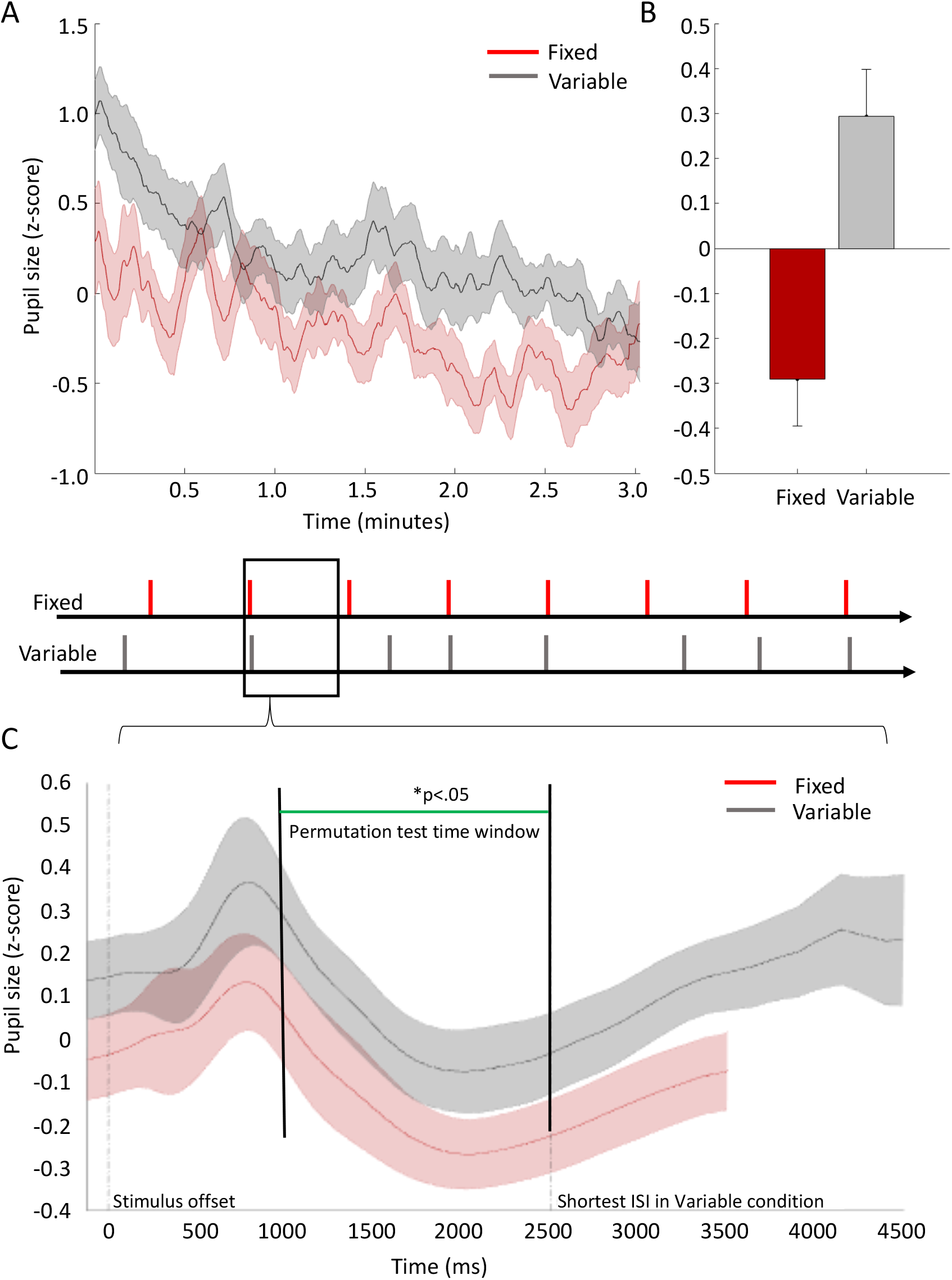
A) average pupil size (standardised) in Fixed blocks (in red) and Variable blocks (in grey) plotted across all participants, over the duration of experimental blocks, lasting approximately 3 minutes. Lines represent the mean and shaded error bars represent standard error. For illustration purposes only, data were smoothed using a time window of 7000 ms (equivalent to an average duration of two trials) and the data were standardised by converting the raw pupil size to y scores. The conversion was done separately for each participant, based on the mean pupil size (and standard deviation) for that individual throughout the entire experimental run (irrespective of experimental condition). (B) contrasting the mean pupil size (standardised z-scores) during Fixed blocks (red bars) vs. Variable blocks (grey bars). Error bars represent standard error. (C) (Standardised) Pupil size over the duration of individual experimental trials averaged across all participants in Fixed blocks (in red) and Variable blocks (in grey). Lines represent the mean and shaded error bars represent the 95% confidence interval range. All trials in Fixed blocks lasted 3500 ms. In Variable blocks, trial duration varied in range between 2500 ms and 4500 ms for the longest trials. Therefore, the grey time series includes a varying and decreasing number of observations within the range of 2500 – 4500 ms following stimulus offset. For illustration purposes only, data were smoothed using a time window of 250 ms (equivalent to the gap between two adjacent interstimulus intervals in the Variable condition). The data were standardised by converting the raw pupil size to z-scores. The conversion was done separately for each participant, based on the mean pupil size (and standard deviation) throughout the entire experimental run (irrespective of experimental condition). We used a permutation t-test to compare the time-series within the range of 1000 to 2500 ms following stimulus offset. The permutation t-test was based on 50,000 samples. The resulting distribution for each data point (each millisecond of recorded data) was compared to a critical t-value (p<.05) adjusted for multiple comparisons based on the ‘t-max’ method (Blair & Karniski, 1993). The green line indicates the time points at which the two time-series differed significantly.

Our second set of analyses characterised anticipation-related short-term changes in pupil diameter. We constructed a time series of the standardised pupil size immediately after stimulus onset and throughout a trial until the onset of the subsequent stimulus, to observe event-locked changes in pupil size. Figure 2c shows the contrast between the mean pupil size in Fixed blocks and in Variable blocks. After the minimal inter-stimulus (2500 ms), the time series in Variable blocks include a varying and decreasing number of observations until the maximum interval of 4500 ms.

We used the stimulus-locked time series to retest for tonic differences, this time at the trial level. We first focused on the full 2500 ms time window immediately after stimulus offset, to include all the trials in both conditions. To compare between conditions, we used a Linear Mixed-effects Model, which included the Block Type and Time as fixed factors, and a maximal random-effects structure (Barr, Levy, Scheepers, & Tily, 2013) that included intercepts for each participant, as well as by-participant slopes for the effects of Block Type and Time. We ran this model using the mean standardised pupil size as the dependent variable (the data depicted in Figure 2c), and identified a significant difference between the two experimental conditions (t(124997)=2.72; *p*=.006; 95%CI[.05;.32]). There was also a main effect of Time (t(124997)=-6.40; *p*<.001; 95%CI[-.0002;-.0001]) but no interaction between Block Type and Time (*p*=.84). A permutation test was also carried out to confirm the results using a statistical approach unaffected by autocorrelations among successive measurements (see: Zokaei, Board, Manohar, & Nobre, 2019, for a similar approach). Permutation t-tests comparing pupil size after the initial stimulus-evoked response (1000 to 2500 ms) showed a consistent difference between the conditions throughout the time window of interest as indicated by the green line in Figure 2c. The permutation test was based on 50,000 iterations and was corrected for multiple comparisons based on the ‘t-max’ method (Blair & Karniski, 1993) (see Methods section).

As seen in Figure 2c, after the initial stimulus-related pupillary response a similar gradual increase in pupil size occurred in both Fixed and Variable conditions, commencing approximately 2000 ms after stimulus onset time. In the Variable condition, the increase in pupil size followed the increasing interval and thus the increasing probability for stimulus presentation. To characterise this trend statistically, we modelled the data using a linear regression. First, for each participant, we calculated the mean time series of pupil size between 2000 ms after onset, until 4500 ms. The data were then fitted and compared to a constant model. The result indicated a significant slope (R^2^=.158; F(1,62498)=11767; p<.001; partial η^2^=.158).

### Experiment 2

#### Behavioural data

Data were modelled separately for each block type (Fixed vs. Variable) using the LibTVA (Dyrholm *et al*., 2011; Kyllingsbaek, 2006), a MATLAB toolbox available online to generate TVA model based on empirical datasets. To validate the reliability of the model, we calculated the correlations between the original data and the predicted data on each exposure duration. The mean Pearson’s r score between the predicted and the original performance data was 0.95, confirming a large effect size. The processing-speed parameter was affected by temporal expectations, with faster processing rate in Fixed (*v*=17_items/minute_; SE=1.6) compared to Variable (*v*=15_items/minute_; SE=1.4) blocks; t(25)=2.18; *p*<.05; 95%CI[.08;3.09]). There was no difference in the perceptual threshold between the two conditions (t(25)=-1.631; *p*=.115; 95%CI[-2.83; .32]).

Comparing the rate of correct responses for targets presented for visible targets (40, 70 or 100 ms; see Methods section) provided a complementary measure of the benefits of temporal expectation on performance. To compare performance in Fixed and Variable blocks, we focused on the common 3500 ms interval (see figure 3b). The results indicated a significant difference in accuracy between conditions (Coefficient Estimate = .09; SE=.04; t(7490)=2.22; p=.025; 95%CI=[.01;.16]). Participants were more accurate when responding to targets presented at a fixed rhythm (.58; SE=.005) compared to variable rhythm (.56; SE=.013). There was also a main effect of exposure duration on accuracy (t(7490)=14.52; p<.001; 95%CI=[.03;.04]) but the two variables did not interact (*p*=.6).

**Figure 3:**
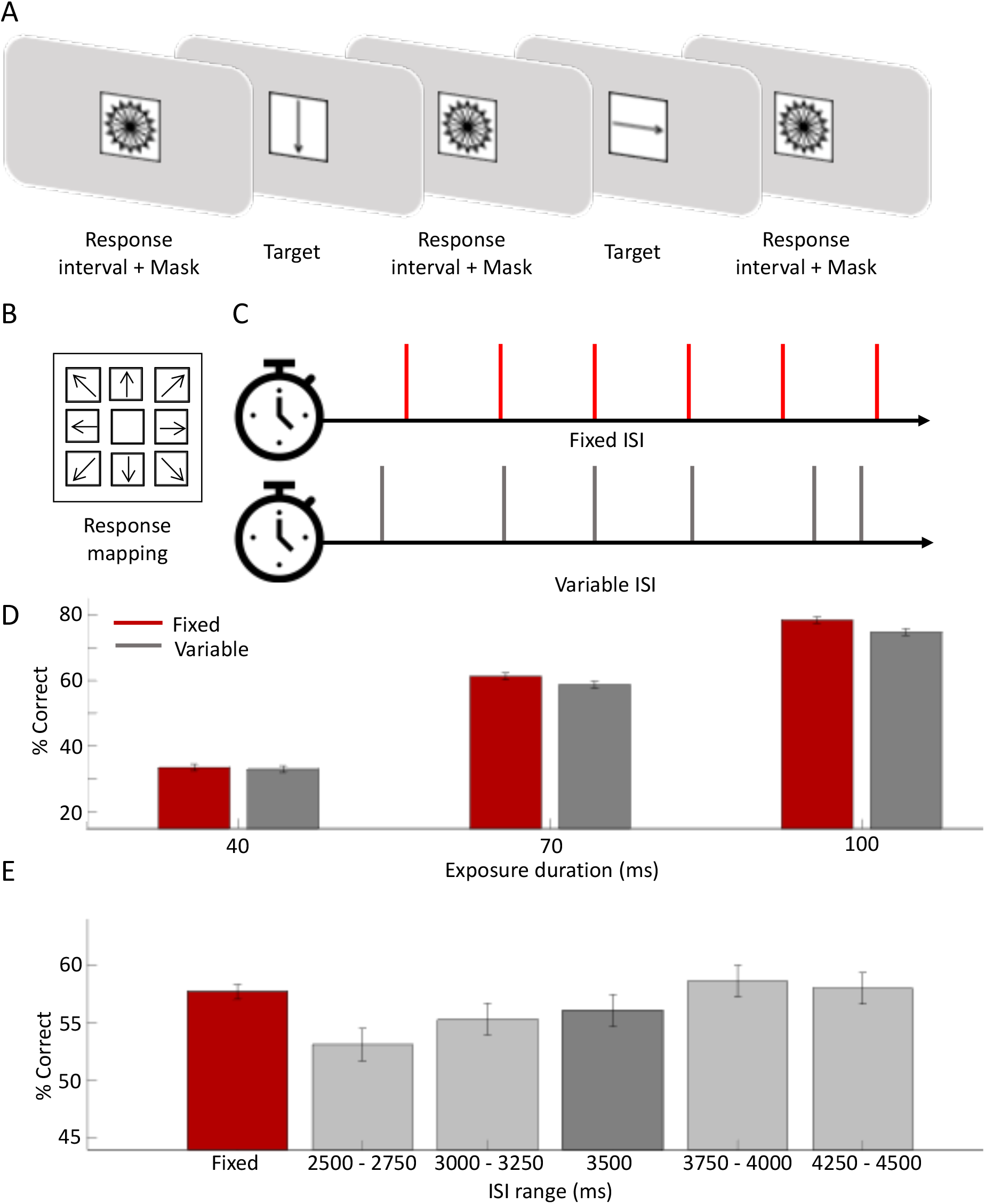
(A) A schematic outline of the experimental design and the response mapping: a visual mask, made of overlapping arrows organized in a circle, appeared at the centre of the screen during the inter-stimulus interval. The inter-stimulus interval was determined according to the experimental condition (with a minimum of 2.5 sec); the mask was replaced by a single arrow, appearing for a variable duration: 10/20/40/ 70/100 ms, and immediately masked again, and participants were instructed to respond while the mask was presented, or do nothing if they did not perceive the target; (B) participants used the keyboard numpad, where eight arrows were drawn on stickers to indicate the response mapping. As in the illustration, the direction of the arrows corresponded to their locations to assist participants and allow them to respond without moving their eyes from the screen. Participants were instructed to respond only if they were fairly certain to have perceived the stimulus; (C) there were two blocked conditions: a fixed block, in which targets appeared in a fixed 3500-ms rhythm (red) and a Variable block in which intervals varied between 2500 and 4500 ms based on a flat distribution of 20 possible intervals equally spaced every 100 ms (D) comparing percentage of correct responses to targets appearing after 3500 ms intervals in the two experimental conditions (Fixed – in red and Variable – in Grey), by exposure duration. Error bars represent the standard error of the mean. (E) The red bar represents overall performance in Fixed blocks, during which all intervals were equal to 3500 ms. The grey bars depict the mean performance on different subsets trials in variable blocks, based on the inter-stimulus intervals. Error bars represent the standard error of the mean

As in Experiment 1, the data in the Variable blocks also revealed benefits in performance that followed the increasing passage of time and hazard rate (figure 3c). To evaluate this pattern statistically, we modelled the data using a GLMM with the ISI range as a fixed factor, and the mean accuracy as the dependant variable (see Methods section for full model specifications). We included only trials in Variable blocks. The result indicated a significant positive coefficient (.084; SE=.025) (t(4450)=3.29; p=.001; 95%CI=[.03;.13]).

#### Pupillometry data

To compare tonic modulation of pupil size between Fixed and Variable blocks, we contrasted the mean pupil size over the full duration of the two different block types. Figure 4a shows the average standardised pupil size throughout entire experimental blocks, separated by block type. The two standardised signals were compared using a paired-sample t-test with the mean standardised pupil size across the entire block on each condition as the dependent variable (see Figure 4b). There was a significant difference (t(25)=-2.94; p=.007; 95%CI[-.58; −.10]), indicating tonically larger pupil sizes in the Variable condition compared to the Fixed condition.

**Figure 4:**
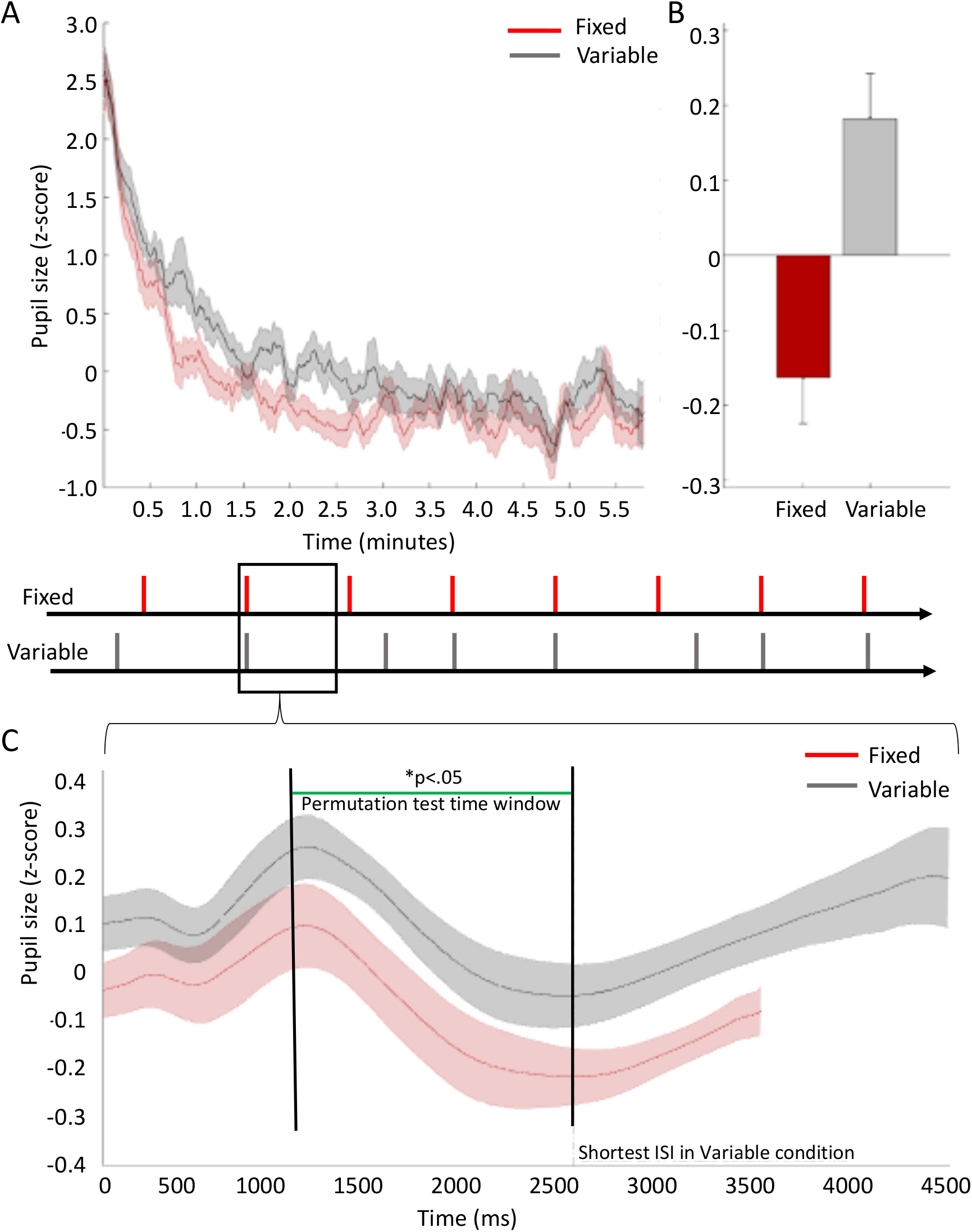
(A) average pupil size (standardised) in Fixed blocks (in red) and Variable blocks (in grey) over the duration of experimental blocks, across all participants, lasting approximately 6 minutes. Lines represent the mean and shaded error bars represent standard error. For illustration purposes, data were smoothed using a time window of 7000 ms (equivalent to an average duration of two trials) and the data were standardised by converting the raw pupil size to z scores. The conversion was done separately for each participant, based on the mean pupil size (and standard deviation) throughout the entire experimental run (irrespective of experimental condition). (B) contrasting the mean pupil size (standardised z-scores) during Fixed blocks vs. Variable blocks. Error bars represent standard error. (C) average pupil size (standardised) in Fixed blocks and Variable blocks over the duration of experimental trials, across all participants. Lines represent the mean and shaded error bars represent the 95% confidence interval range. All trials in Fixed blocks lasted 3500 ms. In Variable blocks, trial duration varied in range between 2500 ms and 4500 ms. Therefore, the grey time series includes a varying number of observations within the range of 2500 – 4500 ms following stimulus offset. For illustration purposes, data were smoothed using a time window of 250 ms. The data were standardised by converting the raw pupil size to z scores. The conversion was done separately for each participant, based on the mean pupil size (and standard deviation) throughout the entire experimental run (irrespective of experimental condition).). We used a permutation t-test to compare the time-series within the range of 1000 to 2500 ms following stimulus offset. The permutation t-test was based on 50,000 samples. The resulting distribution for each data point (each millisecond of recorded data) was compared to a critical t-value (p<.05) adjusted for multiple comparisons based on the ‘t-max’ method (Blair & Karniski, 1993). The green line indicates the time points at which the two time-series differed significantly.

As in Experiment 1, our second set of analyses focused on trial-wise pupil modulation. We constructed time series of the standardised pupil size immediately after stimulus onset and throughout a trial until the onset of the subsequent stimulus, to observe event-locked changes in pupil size. Figure 4c shows the contrast between the mean pupil size in Fixed blocks and in Variable blocks. After the minimal interstimulus interval in Variable blocks (2500 ms), time series include a varying and decreasing number of observations until the maximum interval of 4500 ms.

We used the stimulus-locked time series to retest for tonic differences, this time at a trial level. The approach followed that used for Experiment 1. We focused on the 2500 ms time window immediately after stimulus offset, to include all trials in both conditions. To compare between conditions, we used a Linear Mixed-effects Model, which included the Block Type and Time as fixed factors, and a maximal random-effects structure (Barr et al., 2013) that included intercepts for each participant, as well as by-participant slopes for the effects of Block Type and Time. We ran this model using the mean standardised pupil size as the dependant variable (the data depicted in Figure 4c), and identified a significant difference between the two conditions, indicating relatively larger pupil diameters in the Variable condition (t(129997)=3.03; *p*<.001; 95%CI[.04;.19]). There was no main effect of Time (*p*=.9) or interaction (*p*=.91). A further permutation t-test comparing pupil size after the initial stimulus evoked response (1000 to 2500 ms) confirmed a consistent difference between the conditions throughout the time window of interest, as indicated by a green line on Figure 4c.

As in Experiment 1, a gradual increase in pupil size occurred after the initial stimulus-evoked response in both Fixed and Variable conditions until the onset of the next target. To test for significant increases in pupil size in the Variable condition, we used a linear model to estimate the slope of increase, repeating the same procedure as in Experiment 1: for each participant, we calculated the mean time series of pupil size between 2000 ms after onset, until 4500 ms. The data were then fitted and compared to a constant model. The result indicated a significant positive slope (R^2^=.199; F(1, 64998)=16156; p<.001; partial η^2^=.199).

## General Discussion

Across our two experiments, we found clear evidence that temporal structures embedded in continuous-performance tasks benefit behaviour and modulate both tonic and short-scale pupil dynamics, pointing to efficient regulation of arousal at multiple timescales. When fixed temporal intervals maximised temporal expectations, superior behavioural performance was achieved while a more efficient energetic state was maintained overall. When variable intervals were used, arousal levels were elevated overall but were also sensitive to the passage of time within trials, possibly reflecting conditional probabilities learned over the trial history of the experimental block.

Using pupil size as a proxy measure of arousal, our results revealed striking and reproducible effects of temporal regularities on the arousal function, thus revealing arousal to be flexible and adaptive – adjusting to regularities at different time scales. Within CPT, we observed the tonic level of arousal to adjust in line with the level of temporal uncertainly. The continuous context of our task, in which individuals cannot temporarily disengage between discrete trials, was especially conducive for observing the changing patterns of arousal. In Fixed blocks, with high temporal certainty, arousal levels were lower overall. In Variable blocks, with relatively higher temporal uncertainty, participants maintained tonically elevated levels of arousal. Pupil dilation was also sensitive to local modulations of arousal, following temporal prediction or temporal preparation when intervals were variable. In the context of Variable blocks, we observed transient increases in pupil dilation that followed the passage of time and the mounting temporal conditional probability for the occurrence of relevant event as the trial interval increased between 2500 and 4500 ms.

Interestingly, while our experimental manipulation revealed a direct link between temporal expectations and arousal, as well as temporal expectations and performance – the relationship between pupil size and behaviour was not straightforward. When looking at pupil dynamics in both conditions alongside performance patterns, the relationships seem contingent on the temporal context. For example, performance was highest in a fixed context, when overall pupil size (as a proxy of arousal) was relatively small in comparison the variable condition. However, within a variable context, our data indicates potentially different relationships between pupil size and performance: both performance and pupil diameter increase as a function of time. Whereas our experimental design was not optimised to look at trial-wise correlations, the overall pattern is suggestive of different “modes” of performance resembling other works on “rhythmic” vs. “random” modes (Schroeder, Herrero, & Haegens, 2014) albeit within a different time scale of seconds rather than milliseconds.

Our findings align well with the framework of the Adaptive Gain Theory, which proposes that an increase in uncertainty leads to tonic increases in arousal (Aston-Jones & Cohen, 2005). In the context of reinforcement learning, high levels of uncertainty have been linked to a state of ‘exploration’ and continuous high tonic arousal, whereas reduced uncertainty promotes ‘exploitation’ and reduced tonic arousal (Yu & Dayan, 2005). Whereas previous studies have reported pupillary changes related to un/certainty of stimulus identity and their reward associations (e.g., Friedman, Hakerem, Sutton, & Fleiss, 1973; Lavín, San Martín, & Rosales Jubal, 2014; Preuschoff, ‘t Hart, & Einhauser, 2011; Urai, Braun, & Donner, 2017; Vincent, Parr, Benrimoh, & Friston, 2019), our study extends this notion in sustained-attention context and CPT.

In both Fixed and Variable CPT conditions, there were temporal structures available to aid performance. However, in the Fixed condition these were more robust: anticipation could rely on a high degree of predictability. The high temporal certainty in the Fixed blocks afforded a state of exploitation in which tonic arousal remained low. Participants could anticipate the task-relevant events, and regulate arousal only transiently, to perform at high levels. In contrast, variable blocks required a higher degree of temporal exploration, in which participants ‘forage’ for task-relevant events over time.

In line with previous studies, we were able to reveal changes in pupil size occurring proactively in the period of anticipation before stimulus onset (Friedman et al., 1973; Jennings, Van Der Molen, & Steinhauer, 1998). However, to our knowledge, our study is the first to show the adaptation of arousal operating within continuous-performance tasks. These measures of arousal modulation can be directly linked to temporal expectation, and isolated from motor decisions and reward.

Our behavioural-performance results also provide an important advancement to previous work by showing that temporal expectation improves perceptual capacity within extended task contexts. Although it was previously shown that individuals benefit from temporal anticipation based on rhythmic structures in a continuous task (e.g., Dankner et al., 2017), changes to perceptual accuracy was not measured and behavioural outcomes were restricted to response speed. To our knowledge, we present the first evidence for perceptual benefits of temporal expectation in CPT independent of motor preparation. Our tasks placed no emphasis on motor performance: In Experiment 1, we used a task that yielded individual differences in perceptual parameters by using forward and backward masking (Shalev, De Wandel, Dockree, Demeyere, & Chechlacz, 2017; Shalev et al., 2016; Shalev, Humphreys, & Demeyere, 2018; Shalev, Vangkilde, et al., 2019). In Experiment 2, participants performed fine perceptual discriminations. They were encouraged to focus on accuracy rather than speed and were permitted to skip responses. By manipulating stimulus durations and modelling the results, we replicated the benefit of temporal expectation for visual processing speed previously noted in single-trial tasks (Sørensen et al., 2015; Vangkilde et al., 2013; Vangkilde et al., 2012) in a continuous-performance context.

From a wider perspective, it is important to appreciate that rhythmic CPTs are widely used in basic and clinical research and applications, often employed as an elementary neuropsychological tool when assessing sustained attention. Interestingly, a recent study showed impaired capacity to form temporal predictions on a CPT among individuals diagnosed with ADHD (Dankner et al., 2017). Our findings may therefore carry a further implication: the inability to form temporal predictions may also reflect an inability to make the appropriate adjustments of arousal. Such a hypothesis provides an important link between the potential timing difficulties in ADHD and the capacity of sustaining performance over extended time. Indeed, a significant proportion of popular tasks for cognitive assessment of attention difficulties relies on CPT designs with a fixed rhythmic structure (e.g., Conners & Staff, 2000; Lee & Park, 2006; Robertson et al., 1997). Based on our findings, future inquiries may wish to study sustained performance not only using fixed, but also variable intervals, to understand the interactions between arousal and anticipation among diverse populations.

## Methods: Experiment 1

Experimental procedures were reviewed and approved by the central university research ethics committee of the University of Oxford.

### Participants

Thirty neurotypical young adults participated in this study (20 of whom were female, mean age 25, SD=3.2). The sample size was chosen based on previous and ongoing studies using the same behavioural manipulation (Shalev et al., 2016, 2018; Shalev, Vangkilde, et al., 2019). Pilot data contrasting pupil size in a comparable continuous context (variable and fixed rhythms) have yielded medium to large effect sizes (Cohen’s d values ranging between .5 to .7) using comparable samples (ranging between 25-30 participants per experimental group.) Power was calculated based on an effect size of .6 and a two-tailed significance level of .05 to yield a Power (1-β err probability) of 0.8, and resulted with a minimum sample size of N=24. Participants were recruited through an online research-participation system at the University of Oxford. All were right-handed and had normal or corrected eyesight (based on self-reports). They were compensated for their time (£10 per hour).

### Apparatus

A PC with an i7 processor and a 2-GB video card was used for displaying stimuli and recording behavioural data. The task was generated using Presentation software (Neurobehavioural Systems, Albany, CA). The stimuli were presented on a 24” LED monitor, with a screen resolution of 1080 × 1920 and a refresh rate of 100 Hz. All stimuli were preloaded to memory using the presentation software to minimise temporal variability in stimulus display.

### Experimental design

Participants were requested to respond to target shapes and to ignore distractor shapes (see Figure 1 for a schematic illustration). Targets and distractors briefly (120-ms duration) replaced a multi-shape masking stimulus (mask) that was otherwise continuously present. The masked version of the CPT was originally designed to reveal individual differences using perceptual parameters and has been validated in various populations (Shalev et al., 2016, 2018; Shalev, Vangkilde, et al., 2019). Uniquely, in the current study the temporal regularity with which target and distractor shapes appeared was manipulated across conditions.

The mask comprised four superimposed figures (square, triangle, circle and hexagon) in different colours (blue, red, and green). Its total size was 5 degrees of visual angle, both horizontally and vertically. The mask appeared and remained at the centre of the screen, disappearing only when replaced by either a target or a distractor shape for 120 ms. The mask then reappeared immediately, generating pre- and post-masking of each target or distractor. The target shape was a blue hexagon, and distractor stimuli were either similar in colour to the target (blue circle or diamond), similar in shape (red or green hexagon), or completely different (green circle and red diamond). All distractor types appeared with an equal distribution. Two-thirds of events were targets and the rest were distractors. Distractors and targets appeared at the centre of the screen and occupied a square of 5 degrees of visual angle vertically and horizontally. Participants were instructed to press the ‘space bar’ whenever they detected targets, while ignoring and refraining from responding to distractors, in a continuous stream of stimuli lasting approximately 12 minutes (altogether, 133 target and 67 distractor stimuli appeared in between mask stimuli).

The interstimulus intervals (ISIs) between critical stimulus events (targets and distractors) were manipulated to vary the degree of rhythmic temporal expectation. There were four concatenated blocks of trials (not explicitly signalled) alternating between Fixed and Variable conditions. During Fixed blocks, targets and distractors appeared predictably every 3.5 seconds, in a rhythmic fashion. During Variable blocks, targets and distractors appeared unpredictably at intervals between 2.5 and 4.5 seconds (mean 3.5 seconds) with an equal probability for each ISI (drawn from 9 possible intervals, equally spaced between 2500 ms and 4500 ms in gaps of 250 ms). Fixed and Variable blocks were always interleaved, and their order was randomised among participants.

### Procedure

The experiment took place in a dark testing room. Participants sat 50 cm from the monitor, and a chin rest was used to keep their head still. The experimenter was also sitting in the room, behind the participant, to monitor behaviour. The task instructions appeared on the screen and were explained to the participant by the experimenter, who also answered any questions about the procedure. The session lasted approximately 12 minutes, and commenced with a short practice session of 10 trials.

### Statistical analysis

#### Behavioural benefit

First, we sought to verify whether individuals utilised rhythmic structures to benefit behaviour (as previously observed by (Dankner et al., 2017)). Responses were categorised as being: a correct response (responding to target), a correct rejection (withholding response to distractor), a false alarm (responding to distractor), or an omission (not responding to target). We derived perceptual parameters based on the Signal Detection Theory (SDT; Green and Swets, 1966) to estimate the perceptual sensitivity (d’) and response bias (‘criterion’; β). To correct for extreme cases where performance reached ceiling, we applied a loglinear correction (Hautus, 1995)

First, we focused only on perceptual judgments that followed a 3.5 seconds interval, which was used in the isochronous rhythmic blocks and preceded 20% of the trials in the Variable condition, since this allowed us to learn whether responses to targets were affected by the fixed/variable contexts, while controlling for other temporal factors that may influence performance, such as passage of time or hazard rates (Nobre & Rohenkohl, 2014). Subsequently, to estimate the contribution of passage of time and hazard rates to performance, we calculated the perceptual sensitivity on five ranges of intervals between stimuli: trials that followed 2500 – 2750 ms intervals; 3000 – 3250 ms; 3500 ms; 3750 – 4000 ms; 4250 – 4500 ms. We used an ANOVA test to evaluate a linear contrast, as an indication for an increase in perceptual performance as a function of the interval range.

#### Pupillometry

Pupil size was recorded using a 2000-Hz sampling rate (1000 Hz per pupil), allowing the construction of time series. The analysis procedure first focused on overall differences in pupil size between conditions to identify tonic effects related to temporal expectations *(Tonic Effects* – a further description appears below). Consistent with the task parameters, the optimal time frame for identifying tonic effects within a trial was the period of 2.5 seconds following stimulus onset (2.5 seconds being the shortest ISI and therefore allowing the inclusion of all trials). In addition, we estimated tonic effects by averaging pupil size during entire blocks (50 trials, over 3 minutes). Then, to identify markers of pupil changes due to temporal expectation, we analysed the changes in pupil size during the 1-second period preceding stimulus onset (*Anticipation* – a further description appears below). The raw pupil data were converted to a data matrix using a MATLAB script, excluding data from practice and breaks between blocks. Blinks were interpolated using a cubic spline interpolation (Mathôt et al., 2013), using a dedicated Matlab function available as part of the Pupil Response Estimation Toolbox (PRET) (Denison, Parker, & Carrasco, 2019). Data were smoothed using a moving average widow, based on the ‘smoothdata’ MATLAB function. This method computes a window size based on a heuristic set to attenuate approximately 10% of the energy of the input data (smoothing factor set to 0.1).

Tonic differences in arousal between conditions were calculated based on a direct comparison of standardised pupil size. Raw data were first converted to z scores, by calculating for each participant separately the mean and standard deviation of pupil size throughout the entire experiment and converting all the pupil samples to z scores. We then calculated the mean standardised pupil time-series for each participant, on each block type, which resulted in a time series extending over approximately 3 minutes, consisting ~180,000 samples (sampling rate in milliseconds). We then compared the overall mean standardised pupil size for each participant on each condition using a repeated-measures permutation t-test (based on 50,000 repeated samples).

To look at trial-level modulation of pupil diameter, we constructed two timeseries of different lengths: 3500 ms in Fixed blocks, which was the same interval throughout the experiment; and 4500 ms, which was the maximum length of a single trial in Variable blocks. After the minimal inter-stimulus interval in Variable blocks (2500 ms), time series include a varying and decreasing number of observations until the maximum interval of 4500 ms. These data were used in multiple ways.

First, we used LMM to revalidate the presence of tonic differences in pupil size also when inspecting single trials. We modelled the standardised pupil size as the dependant variable, over 2500 ms to include all the trials in both conditions. We included the Block Type and Time as fixed factors, and a maximal random-effects structure (Barr et al., 2013) that included intercepts for each participant, as well as by-participant slopes for the effects of Block Type and Time. These tonic differences were then confirmed using a more focused approach, contrasting the two time-series directly with a permutation t-test based on 50,000 samples. The resulting distribution for each data point (each millisecond of recorded data) was compared to a critical t-value (p<.05) adjusted for multiple comparisons based on the ‘t-max’ method (Blair & Karniski, 1993). This approach was used by previous studies to assess changes in pupil diameter over time (e.g., (Zokaei et al., 2019)). We focused on a time window between 1000 ms and 2500 ms after the onset of the visual mask which immediately followed a target or distractor stimuli. This time window was chosen to reduce the influence of the transient pupil responses caused by a change in visual stimulation and matches previous work in our lab (e.g., (Zokaei et al., 2019)) as well as our pilot data (see: (Shalev, 2017; Shalev, Demeyere, & Nobre, 2017; Shalev & Nobre, 2018))

Next, we tested for effects of temporal preparation or anticipation based on the passage of time and mounting conditional probabilities at the single-trial level. We focused on the time series in Variable blocks, which included multiple interstimulus intervals. We used a linear regression to evaluate the slope of change in pupil size between 2000 ms after onset until 4500 ms, based on the mean pupil time-series we constructed for each participant. The data were then fitted and compared to a constant model.

#### Data exclusion

Before running statistical comparisons, we evaluated the number of usable trials for each participant. Usable trials were those in which there was a valid record of the pupil in at least one eye (left or right), and in which eyes were fixated at the centre of the screen. As a default, we focused on analysing the data from the right eye. We identified five participants who had a substantial number of missing trials (50% and more), and they were excluded from the study prior to any further analysis. The final set of analyses included 25 participants.

## Methods: Experiment 2

To test for the reliability and generalisability of behavioural benefits and arousal modulation by temporal expectations in continuous performance tasks, we used a task that required discrimination of masked targets appearing either in a fixed rhythm or at variable intervals. The task parameters were designed to allow modelling of the behavioural benefits of predictability using the Theory of Visual Attention (TVA) (Bundesen, 1990). Previous TVA modelling in studies of temporal expectation using discrete trials has revealed performance benefits linked to increases in the parameter modelling Visual Processing Speed (*v*) (Sørensen, Vangkilde, & Bundesen, 2015; Vangkilde, Petersen, & Bundesen, 2013; Signe Vangkilde, Coull, & Bundesen, 2012). We tested whether these findings extended to a continuous-performance context and then turned to our main question about whether tonic and phasic modulations of pupil size similarly accompanied effects of temporal expectation as in our first experiment.

### Participants

Participants in this experiment were 30 naïve volunteers (20 of whom were female, mean age 24.5, SD=4.18). They were recruited through an online research-participation system at the University of Oxford. All had normal or corrected eyesight. Five were left-handed and the rest were right-handed (based on self-reports). They were compensated for their time (£10 per hour).

### Apparatus

A PC with an i7 processor and a 2-GB video card was used for displaying stimuli and recording behavioural data. The task was generated using Presentation software (Neurobehavioural Systems, Albany, CA). The stimuli were presented on a 24” LED monitor, with a screen resolution of 1080 × 1920 and a refresh rate of 100 Hz. All stimuli were preloaded to memory using the presentation software to guarantee minimal temporal noise. A video-based eye-tracker (EyeLink 1000, SR Research, Ontario, Canada) was used to measure pupil diameter as well as to monitor eye movements and blinks at 1000 Hz. The recorded data were saved to an eye-tracking PC.

### Stimuli

Participants in Experiment 2 performed a continuous task in which they had to make perceptual discriminations based on stimuli that briefly interrupted a mask at regular (3500 ms, Fixed condition) or variable (2500 – 4500 ms, Variable condition) intervals. In order to apply TVA to test the nature of benefits conferred by temporal expectation in this CPT, the task required a finer perceptual discrimination on stimuli that varied in duration across trials. Participants were asked to report the direction of arrow stimuli that occasionally interrupted a mask stimulus during extended task blocks (see Figure 3a for a schematic). The mask was a black stimulus comprising 16 bi-directional black arrows circumscribed in a black circle appearing at the centre of the screen (see Figure 3a). The total size of the mask was approximately 4.5° of visual angle horizontally and vertically. Following the presentation of the mask, a single target arrow pointing at one of eight possible directions (in a square occupying 4.5°) appeared for a varying duration (10/20/40/70/100 ms) and was immediately replaced by the mask. Participants were instructed to try to identify the target arrow and to indicate its direction using the arrow numpad, in which eight arrows pointing at different directions appeared at a corresponding location (see Figure 3a). Responses were recorded during the successive inter-stimulus interval, during which the mask was presented.

The task consisted of 8 blocks with 100 trials and each lasting approximately 6 min. The blocks alternated between two conditions, which varied according to temporal expectation. On a Fixed block, a target appeared predictably every 3500 ms; on a Variable block, the target appeared unpredictably at intervals between 2500 ms and 4500 ms (mean 3500 ms) with an equal probability for each ISI (drawn from 20 possible intervals, equally spaced between 2500 ms and 4500 ms). Fixed and Variable blocks were always interleaved. Half the participants commenced with a Fixed block, and the other half with a Variable block.

We estimated how well participants identified ‘targets’ (arrow figures) that appeared at the centre of the screen every few seconds, and compared performance when the onset of the targets was temporally predictable vs. unpredictable using a computational model based on the theory of visual attention (Bundesen, 1990; Dyrholm, Kyllingsbæk, Espeseth, & Bundesen, 2011). We also compared simple accuracy measures for visible targets (40, 70, and 100-ms exposure) between conditions.

### Procedure

The experiment was conducted in a dark testing room. Participants sat 50 cm from the monitor, and a chin rest was used to keep their head still. The eye-tracking device was placed near the monitor and was set to record the two eyes by default. The session began with a short procedure of calibrating the eye tracker. The task instructions then appeared on the screen and were explained to the participant by the experimenter. The participants were told that a visual mask would appear at the centre of the screen, to be replaced briefly every few seconds by a single arrow pointing in one of eight different directions, corresponding to the arrows on the numpad. They were also told that the arrow would be replaced immediately by a visual mask. Whenever the mask appeared, they were to indicate the direction in which the target arrow pointed.

Participants were asked to be as accurate as possible and to keep their eyes fixed on the mask while responding. The analogous spatial organisation of the response mapping on the numpad made it possible to respond while maintaining fixation. It was also emphasised that the speed of responses was not important. Participants were told that if they did not perceive any arrow, they could simply skip the trial. However, they were encouraged to guess if they were ‘fairly certain’ of having identified the arrow, in line with instructions in previous TVA studies (Sørensen et al., 2015). At the end of each experimental block, participants received feedback indicating their accuracy level. Accuracy was calculated based only on the responses provided, ignoring skipped trials. Participants were asked to aim for 80%-90% accuracy and thus encouraged to guess less if their accuracy fell below 80% and to guess more if performance exceeded 90%.

Before beginning the experimental session, a short practice session with variable inter-stimulus intervals (2500 – 4500 ms) consisting of 20 trials allowed participants to learn the task. During the practice block, the first 10 trials presented the targets for an extended duration of 120 ms. The experimenter monitored the participants’ responses at this stage to ensure they understood and followed instructions. The experiment itself consisted of eight blocks (each experimental block lasting approximately 6 minutes). Each block was followed by a short break (approximately one minute). In each block there were 100 trials. There were four Fixed blocks and four Variable blocks. In all, there were 800 experimental trials (400 for each condition) excluding practice.

### Statistical Analysis

#### Behavioural data

We tested whether temporal predictions influenced performance parameters in a continuous-performance task using TVA modelling parameters and simple accuracy measures.

For the modelling analysis, the data extraction and fitting procedures were performed using Matlab (Ver. 2019a; MathWorks, Inc., Natick, MA) and the LibTVA (Dyrholm et al., 2011; Kyllingsbæk, 2006). The behavioural dataset was first split according to experimental condition: Fixed (4 blocks) or Variable (4 blocks). The calculation of the theoretical attentional parameters (processing speed and perceptual threshold) was based on a maximum-likelihood fitting procedure introduced by Kyllingsbaek (2006) to model the observations based on the TVA framework, using the default modelling parameters (K model ‘free’; exponential fitting). The fitting algorithm output includes two theoretical parameters: (1) Parameter *to* is the perceptual threshold, defined as the longest exposure duration that does not evoke conscious perception, measured in seconds; (2) Parameter *v* is the visual processing speed, or processing rate, measured in the number of target items processed per second. We then compared the resulting parameters between the two conditions.

For the analysis of the empirical data, we compared the accuracy rate between conditions for targets presented for 40, 70 and 100 ms. We focused on these exposure durations, as eight participants had missing valid responses in shorter durations. We used a Generalised Linear Mixed-Effect (GLME) model for this comparison, to compare target identification in two conditions (comparable to our contrast in Experiment 1) while also accounting for the additive effect of exposure duration on target detection. Thus, the model included the effects of Block Type (Variable vs. Fixed) and Exposure Duration (40, 70, and 100 ms) as fixed factors, and a random-effects structure that included intercepts for each participant, as well as by-participant slopes for Block Type and Exposure Duration.

#### Pupillometry data

we repeated the same procedure as in Experiment 1.

#### Data exclusion

We applied the same criteria as in Experiment 1. Four participants had a substantial number of missing trials (50% and more) and were excluded prior to any further analysis. The final set of analyses included 26 participants.

## Acknowledgments

This work was supported by a Wellcome Trust Senior Investigator Award (104571/Z/14/Z) and a James S. McDonnell Foundation Understanding Human Cognition Collaborative Award (220020448) to A.C.N, and by the NIHR Oxford Health Biomedical Research Centre. The Wellcome Centre for Integrative Neuroimaging is supported by core funding from the Wellcome Trust (203139/Z/16/Z).

## Author Contributions

NS and ACN developed the study concept and wrote the manuscript. NS programmed the experiment, collected, and analysed the data.

